# Dissecting the herpesvirus architecture by targeted proteolysis

**DOI:** 10.1101/311894

**Authors:** Gina R. Daniel, Caitlin E. Pegg, Gregory A. Smith

## Abstract

Herpesvirus particles have a complex architecture consisting of an icosahedral capsid that is surrounded by a lipid envelope. Connecting these two components is a layer of tegument that consists of varying amounts of twenty or more proteins. The arrangement of proteins within the tegument cannot easily be assessed and instead is inferred from tegument interactions identified in reductionist models. To better understand the tegument architecture, we have developed an approach to probe capsid-tegument interactions within extracellular viral particles by encoding tobacco etch virus (TEV) protease sites in viral structural proteins, along with distinct fluorescent tags in capsid and tegument components. In this study, TEV sites were engineered within the pUL36 large tegument protein: a critical structural element that is anchored directly on the capsid surface. Purified pseudorabies virus extracellular particles were permeabilized and TEV protease was added to selectively cleave the exposed pUL36 backbone. Interactions with the capsid were assessed in situ by monitoring the fate of the fluorescent signals following cleavage. Although several regions of pUL36 are proposed to bind capsids, pUL36 was found stably anchored to the capsid exclusively at its carboxyl terminus. Two additional tegument proteins, pUL37 and pUS3, were tethered to the capsid via pUL36 whereas the pUL16, pUL47, pUL48, and pUL49 tegument proteins were not stably bound to the capsid.

**IMPORTANCE:** Neuroinvasive alphaherpesviruses produce diseases of clinical and economic significance in humans and veterinary animals, but are predominantly associated with less serious recurrent disease. Like all viruses, herpesviruses assemble a metastable particle that selectively dismantles during initial infection. This process is made more complex by the presence of a tegument layer that resides between the capsid surface and envelope. Components of the tegument are essential for particle assembly and also serve as critical effectors that promote infection upon entry into cells. How this dynamic network of protein interactions is arranged within virions is largely unknown. We present a molecular approach to dissect the tegument and with it, begin to tease apart the protein interactions that underlie this complex layer of the virion architecture.

## INTRODUCTION

Viruses from the order *Herpesvirales* infect animals ranging from invertebrates to humans, with over 300 herpesviruses identified to date (1). Despite their wide prevalence, these viruses share a common particle morphology and the ability to persist throughout the lifetime of their hosts (2). Neurotropic members of the *Alphaherpesvirinae* subfamily of the *Herpesviridae* family include clinical (e.g. herpes simplex virus type I; HSV-1) and veterinary (e.g. pseudorabies virus; PRV) viruses that infect mucosal-epithelial surfaces before transmitting to innervating neurons of the peripheral nervous system (PNS) (3).

Herpes virions are ~225 nm in diameter and consist of a linear double-stranded DNA genome, an icosahedral protein capsid, a protein tegument layer, and a lipid envelope containing viral glycoproteins (2, 4). Virion production begins in the nucleus, where capsid assembly and genome encapsidation occurs, and concludes in the cytoplasm with the envelopment of the capsid-tegument complex (5). While the structure of the capsid is solved to 3.1 Å for HSV-1 and 4.9 Å for PRV (6–8), the majority of the tegument layer departs from the radial symmetry of the capsid shell and cannot be visualized in great detail by composite averaging methods such as cryo-electron microscopy (9). This technical limitation is overcome by the application of cryo-electron tomography, but this method has yet to achieve resolutions necessary to infer protein distributions within the herpesvirus virion (10, 11). Despite this lack of architectural information, the constituents of the tegument layer are defined along with many pairwise interactions between them (2, 5, 12, 13). In addition, single-particle fluorescence analysis has revealed tegument protein localization to specific sub-regions of the viral particle (14, 15). Despite these advances, the question of how the tegument is organized remains poorly defined. Most of the inner tegument density is localized to capsid vertices, and to date, three tegument proteins have been proposed to contact the capsid directly: pUL16, pUL36, and pUL47 (16–19). The best established capsid-tegument interaction is the pUL36 protein, which is directly anchored to the pUL25 capsid protein as part of a five-helix bundle around capsid vertices (6, 7, 18, 20–22).

The pUL36 protein is a large (330 kDa in PRV) and conserved tegument constituent (5). Since the initial finding that the C-terminus of pUL36 binding to capsids via pUL25 in HSV-1 and PRV (23), other regions of this large tegument protein have also been proposed to bind the capsid (21, 24–26). In this study, we sought to identify how pUL36 and other tegument proteins interact with the capsid in PRV. Because pUL36 is essential for viral replication, it was necessary to use an approach that was conditional and did not disrupt the formation of mature viral particles (27–33). To this end, we developed a selective on-particle disassembly assay to investigate capsid-tegument interactions in virions. We demonstrate that the C-terminus of pUL36 is solely responsible for its stable attachment to capsids, and that capsid associations with tegument proteins pUS3 and pUL37 are dependent on pUL36. No evidence of additional capsid-tegument interactions was obtained under the conditions employed in this analysis.

## MATERIALS AND METHODS

### Cells and Virus Strains

Recombinant PRV strains used in this study were constructed by two-step, RED-mediated recombination with the pBecker3 infectious clone (34, 35). A summary of the recombinant strains of PRV used in this study is provided in Table 1, and the primers used for their production are listed in Table 2. Tobacco etch virus (TEV) protease recognition sites and the GFP coding sequence were inserted into the pBecker3 derivative pGS4284, which encodes a pUL25/mCherry (red-fluorescent capsid) fusion, and pGS5507, which additionally encodes a GFP-pUL36 fusion (36, 37). Recombinant clones were initially identified by restriction enzyme analysis and confirmed by sequencing. Infectious clones were transfected into PK15 epithelial cells as previously described to produce stocks of recombinant PRV (38). Working stocks were made by an additional passage through PK15 cells. PK15 cells were maintained in Dulbecco’s modified Eagle medium (DMEM) supplemented with 10% (vol/vol) bovine growth supplement (BGS; HyClone). During infection, the BGS concentration was reduced to 2%. Virus stocks were titered on PK15 cells as previously described (39). Titer measurements were compiled from a minimum of 3 individual working stocks. Individual plaque diameters on PK15 cells were measured from the average of two orthogonal cross-sections based on pUL25/mCherry fluorescence emissions. Greater than 30 plaques were measured per experiment. This process was repeated three times to arrive at the average plaque size for each virus, which was normalized as a percent of the parental virus plaque diameter.

**Table 1:** Recombinant PRV used in this study

| Strain | Capsid tag | Tegument tag | TEV insertion site in pUL36 | Titer (PFU/ml) | Source |
| --- | --- | --- | --- | --- | --- |
| PRV-GS4848 | mRFP1-GFP-VP26 | - | - | 0.2 x 10<sup>9</sup> | Bohannon, 2013 |
| PRV-GS4284 | UL25/mCherry | - | - | 1.0 x 10<sup>9</sup> | Bohannon, 2012 |
| PRV-GS5507 | UL25/mCherry | GFP-pUL36 | - | 1.5 x 10<sup>9</sup> | Huffmaster, 2015 |
| PRV-GS5962 | UL25/mCherry | GFP-pUL36 | aa317/318 (A) | 1.2 x 10<sup>9</sup> | This study |
| PRV-GS6046 | UL25/mCherry | GFP-pUL36 | aa482/483 (B) | 1.4 x 10<sup>9</sup> | This study |
| PRV-GS6047 | UL25/mCherry | GFP-pUL36 | aa599/600 | 0.7 x 10<sup>9</sup> | This study |
| PRV-GS6048 | UL25/mCherry | GFP-pUL36 | aa2021/2022 | 0.4 x 10<sup>9</sup> | This study |
| PRV-GS6210 | UL25/mCherry | GFP-pUL36 | aa2622/2623 (C) | 1.6 x 10<sup>9</sup> | This study |
| PRV-GS6220 | UL25/mCherry | - | aa2622/2623 (C) | 2.4 x 10<sup>9</sup> | This study |
| PRV-GS6325 | UL25/mCherry | pUL36-GFP | aa2622/2623 (C) | 0.9 x 10<sup>9</sup> | This study |
| PRV-GS6267 | UL25/mCherry | pUL37-GFP | aa2622/2623 (C) | 0.5 x 10<sup>9</sup> | This study |
| PRV-GS6269 | UL25/mCherry | pUL49-GFP | aa2622/2623 (C) | 0.7 x 10<sup>9</sup> | This study |
| PRV-GS6292 | UL25/mCherry | pUL48-GFP | aa2622/2623 (C) | 0.7 x 10<sup>9</sup> | This study |
| PRV-GS6294 | UL25/mCherry | pUS3-GFP | aa2622/2623 (C) | 1.7 x 10<sup>9</sup> | This study |
| PRV-GS6372 | UL25/mCherry | pUL16-GFP | aa2622/2623 (C) | 1.1 x 10<sup>9</sup> | This study |
| PRV-GS6374 | UL25/mCherry | pUL47-GFP | aa2622/2623 (C) | 1.2 x 10<sup>9</sup> | This study |

**Table 2:**
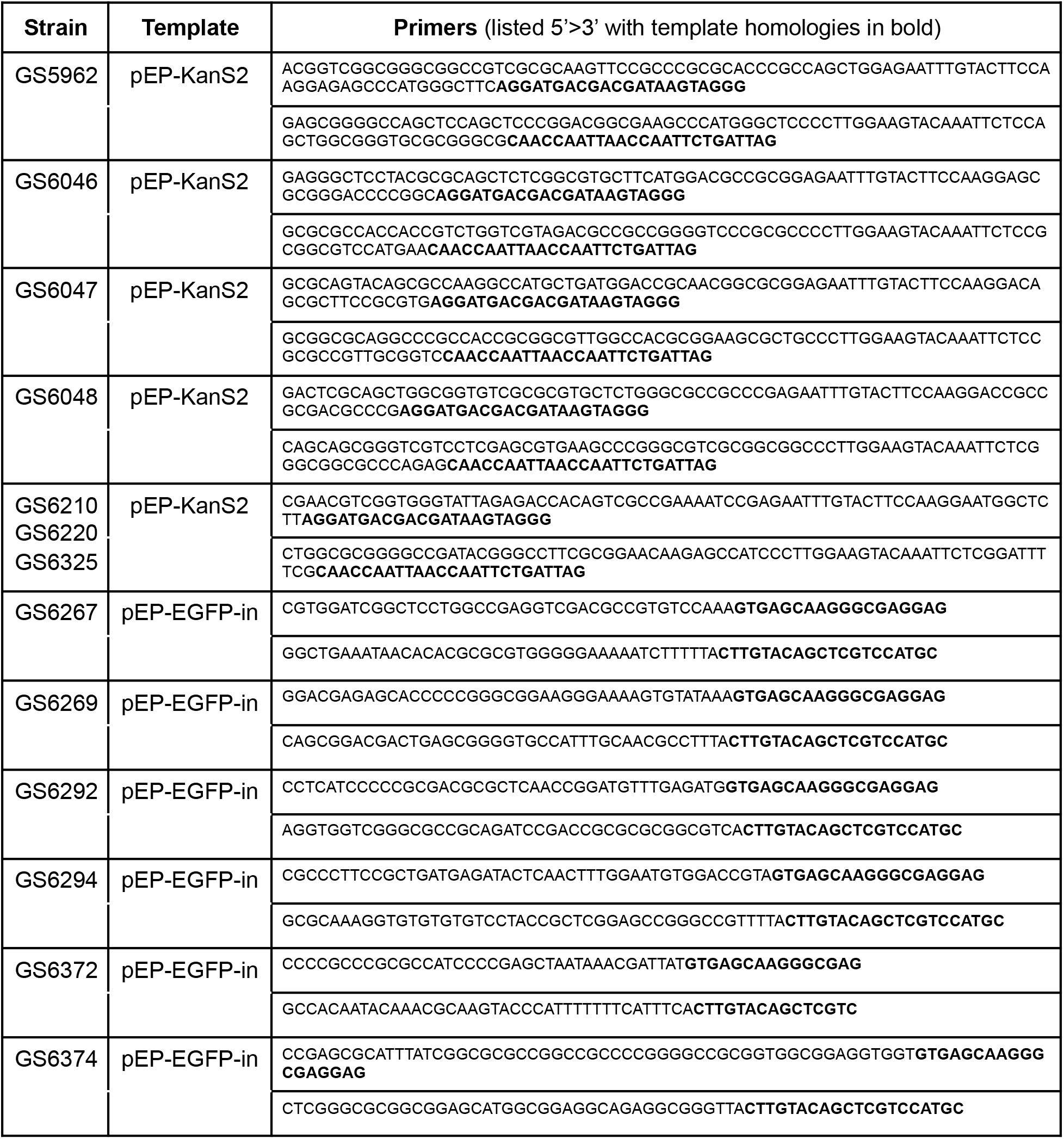
Primers used to construct PRV recombinants unique to this study

### Western blot analysis

PK15 cells were infected at a MOI of 10 with recombinant PRV encoding GFP-pUL36 fusions or GFP fusions to other tegument proteins (Table 1). 5x final sample buffer (FSB: 60 mM Tris [pH 6.8], 10% [vol/vol] glycerol, 0.01% bromophenol blue, 2% [wt/vol] SDS, 5% [vol/vol] β-mercaptoethanol) was added to either purified extracellular virions or extracted & TEV-treated particles. Samples were boiled for 5 min, and separated on 4-20% SDS polyacrylamide gradient (Mini-PROTEAN TGX gel, BioRAD). Proteins were transferred to 0.45 μm pore size Immobilon-FL membranes (Millipore), and blocked in 5% (wt/vol) powdered milk in PBS (13.7 mM NaCl, 0.27 mM KCl, 1 mM Na2HPO4, 0.2 mM KH2PO4). PRV pUL36 was detected using a previously described anti-peptide rabbit antibody at a dilution of 1:2500 (40). PRV VP5 was detected using the mouse monoclonal anti-VP5 clone 3C10 (a gift from Lynn Enquist, Princeton University) at a dilution of 1:1000. The mouse monoclonal GFP clone B-2 (Santa Cruz, sc-9996) was used to detect GFP-tagged proteins at a dilution of 1:1000. Antibodies were prepared in 1% milk-PBST (13.7 mM NaCl, 0.27 mM KCl, 1 mM Na2HPO4, 0.2 mM KH2PO4, 0.1% Tween-20). Detection was performed using a LI-COR Odyssey Fc imaging system with donkey anti-rabbit IRDye 680RD or goat anti-mouse IRDye 800CW secondary antibodies at a dilution of 1:10,000 (LI-COR Biosciences).

### Preparation of extracted viral particles

Approximately 4.6 x 10^7^ cells in six 15 cm plates were infected with PRV strains at an MOI of 10 PFU/cell and incubated for 24 h in DMEM media (Gibco) supplemented with 2% (vol/vol) BGS. Supernatants (approximately 15 ml per plate) were cleared of cell debris by a 20 min, 5000 × g fixed-angle centrifugation in two 50 ml conical tubes at 4°C. The cleared supernatant was split to three SW28 centrifugation tubes, and each was underlaid with 3 ml of 10% (wt/vol) Nycodenz in TNE buffer (20 mM Tris [pH 7.6], 500 mM NaCl, 1 mM EDTA). Viral particles were pelleted through the Nycodenz cushions by centrifuging for 1 h at 13,000 rpm at 4°C in a SW28 rotor. The pellets were resuspended and combined in TNE buffer (20 mM Tris [pH 7.6], 500 mM NaCl, 1 mM EDTA) and dispersed by gentle pipetting, resulting in the intact viral particle preparation. In some instances the viral particle preparation was extracted in 1 ml of buffer containing 2% Triton X-100 (TX-100) in TNE supplemented with 2 μl of protease inhibitor cocktail (Sigma P8340) for 20 min on ice. The extracted particles were separated from solubilized membrane and proteins by diluting the 1 ml preparation with 4 ml TNE, overlaying onto a 5 ml 35% sucrose cushion in TNE buffer, and centrifuging for 1 hr at 25,000 rpm in a SW41 rotor at 4°C. The pellet was resuspended in PBS and dispersed by gentle pipetting, resulting in the extracted viral particle preparation. For light microscopy, resuspended intact or extracted particles were diluted with PBS and imaged on a plasma-treated No. 1.5, 22 × 22 mm coverslip in a wax-sealed microscope slide chamber as previously described (14, 41).

### TEV treatment of extracted viral particles

Approximately 2 x 10^6^ extracted viral particles in PBS were incubated in a ProTEV Plus (Promega) reaction mix containing 1x ProTEV buffer (50 mM HEPES [pH 7.0], 0.5 mM EDTA) supplemented with 1 mM DTT and either 5-10 U of ProTEV or an equal volume of ProTEV storage buffer lacking enzyme (50 mM HEPES [pH 7.5], 300 mM NaCl, 1 mM DTT, 1 mM EDTA, 50% glycerol, 0.1% Triton X-100) for 18 hours at 30°C in the dark. Following the incubation, particles were diluted in 5x FSB for western blot analysis or were diluted and dispersed in PBS for imaging by fluorescence microscopy.

### Quantitative fluorescence microscopy

Particles were imaged on an inverted wide-field Nikon Eclipse TE2000-E microscope fitted with a 60x 1.4NA objective, a Cascade II:512 camera, and a Lumen PRO Fluorescence Illumination System set at 100% light output. mRFP1 and mCherry fluorescence were imaged using a 572/35 nm excitation filter and a 632/60 nm emission filter; eGFP was imaged using a 470/40 nm excitation filter and a 525/50 nm emission filter (Chroma Technology Corporation). All fluorophores were imaged with the same dual-band-pass dichroic mirror. With one exception, red and green emissions were sequentially captured using 1.7 second exposures, with 1 MHz digitizer and zero electron-multiplying gain. The exception was images captured of PRV-GS6374 particles, which had green emissions approached saturation and were therefore captured with 1.0 second exposures that were then linearly scaled by a factor of 1.7. Fluorescent intensities from individual particles were enumerated with an automated, custom algorithm as described previously (14). Briefly, the algorithm identified diffraction-limited red-fluorescent punctae consistent with individual capsids. Red-fluorescent capsids outside of ±3 standard deviations were post filtered from the data sets to remove noise and virion clusters. Red and spatially-correlated green intensity measurements from the extracellular viral particles were compiled from three independent experiments, with 800-1,300 particles measured per experiment. Nonlinear Gaussian regression was performed in GraphPad Prism. Red intensity measurements were converted to protein copy number by normalizing to the reported copy number for PRV pUL25/GFP (70.5 copies/virion) and green intensity measurements were converted to copy numbers by normalizing to the reported copy number for PRV GFP-pUL36 (107.1 copies/virion) (14). Protein copy numbers were plotted as bar graphs from three independent experiments (two-way ANOVA, with a multiple comparison’s test).

## RESULTS

### Precision cleavage of pUL36 in viral particles

The large tegument protein, pUL36, is a conserved virion component that is directly bound to the capsid surface (21, 23) and is essential for virus propagation (39, 42, 43). Although several mutagenesis studies have provided insights into pUL36 function (28, 29, 31, 44–48), the role of pUL36 in virion assembly has largely been refractory to investigation because of its presumably essential role and the challenge of avoiding spontaneous genetic repair when trans-complementing mutant viruses with the 9 kb UL36 open reading frame. Therefore, to dissect the PRV architecture, tobacco etch virus (TEV) protease cleavage sites were introduced into the pUL36 tegument protein. TEV protease recognizes a seven amino acid sequence, ENLYFQG, that is commonly included in recombinant proteins to allow for proteolytic removal of affinity tags following purification (49). The TEV protease recognition sequence is not predicted to be present in any PRV proteins based on the viral genomic sequence (50). Recombinant PRV strains were produced encoding red-fluorescent capsids (pUL25/mCherry), green-fluorescent pUL36, and a single TEV protease site within pUL36 (Fig. 1A) (36). Regions of pUL36 that exhibit low conservation were chosen as sites of TEV insertion, with the rationale that these areas would be tolerant of the small insertions and accessible for cleavage in the mature protein. The recombinant viruses produced plaques and titers that approximated the wild type (Fig. 1B and Table 1). Because pUL36 is critical for virion assembly and infectivity, these results indicated that TEV-site insertion did not substantially impair pUL36 function (39, 42, 43).

**Figure 1:**
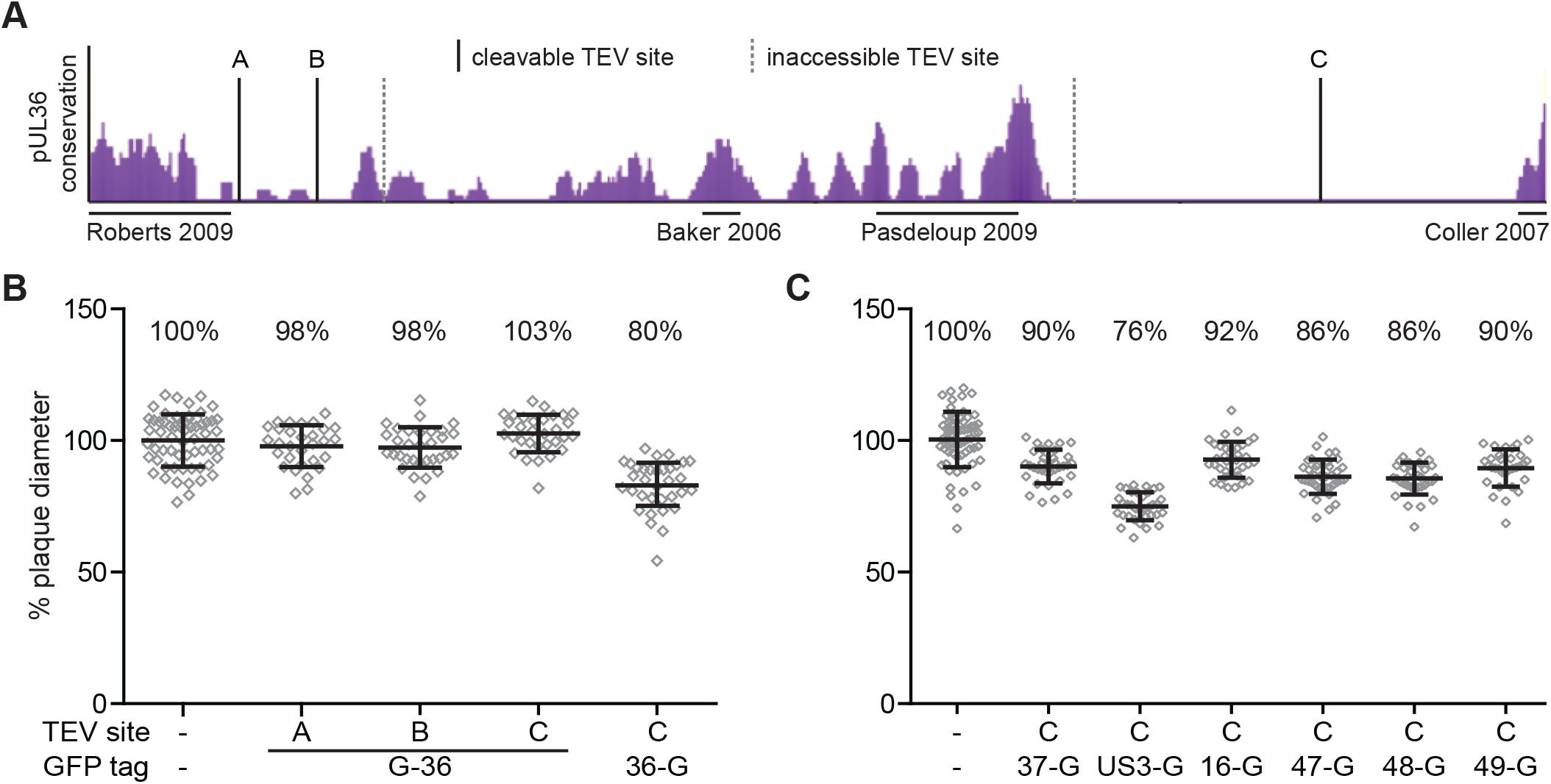
Recombinant PRV strains used in this study. (A) The degree of conservation across the pUL36 predicted amino acid sequence from seven alphaherpesviruses is shown as purple peaks (eShadow). The locations of TEV site insertions are indicated by vertical lines: solid lines were cleavable with exogenous TEV protease and dotted lines were refractory to cleavage - see text). Regions of pUL36 suggested or shown to interact with capsids are marked by horizontal lines under the conservation plot. (B & C) Plaque diameters of recombinant viruses encoding a pUL25/mCherry capsid fusion, a GFP fusion to a designated tegument protein, and a TEV site within pUL36 were compared to a parental virus containing only the pUL25/mCherry tag. Diameters are presented as % of the parental virus.

Virions of each recombinant virus were collected from the supernatant of infected PK15 cells at 24 hours post-infection. After a clarifying step to remove cell debris, viral particles were pelleted from the supernatant through a Nycodenz cushion by ultracentrifugation. The envelopes of the recovered particles were extracted using nonionic detergent (2% TX-100) following a previously described procedure (20). The particles were then pelleted through a 35% sucrose cushion to remove detergent and solubilized material and were incubated in a reaction mixture with or without TEV protease (Fig. 2A). Treatment with TEV protease resulted in cleavage of pUL36 when the TEV site was placed at position A, B or C, as determined by western blot analysis (Fig. 2B). The N-terminal fragments of pUL36 that were produced were consistent with molecular weight predictions based on the locations of these sites. TEV protease did not cleave pUL36 in the remaining two viruses, suggesting the TEV sites at these positions were inaccessible to the protease (Fig. 1A, dashed vertical lines). Importantly, pUL36 cleavage was not observed when TEV protease was omitted (mock treatment) or in a virus that lacked a TEV site (Fig. 2B). These results demonstrated that TEV cleavage was specific and that cleavage occurred efficiently at TEV sites in position A, B, or C within pUL36.

**Figure 2:**
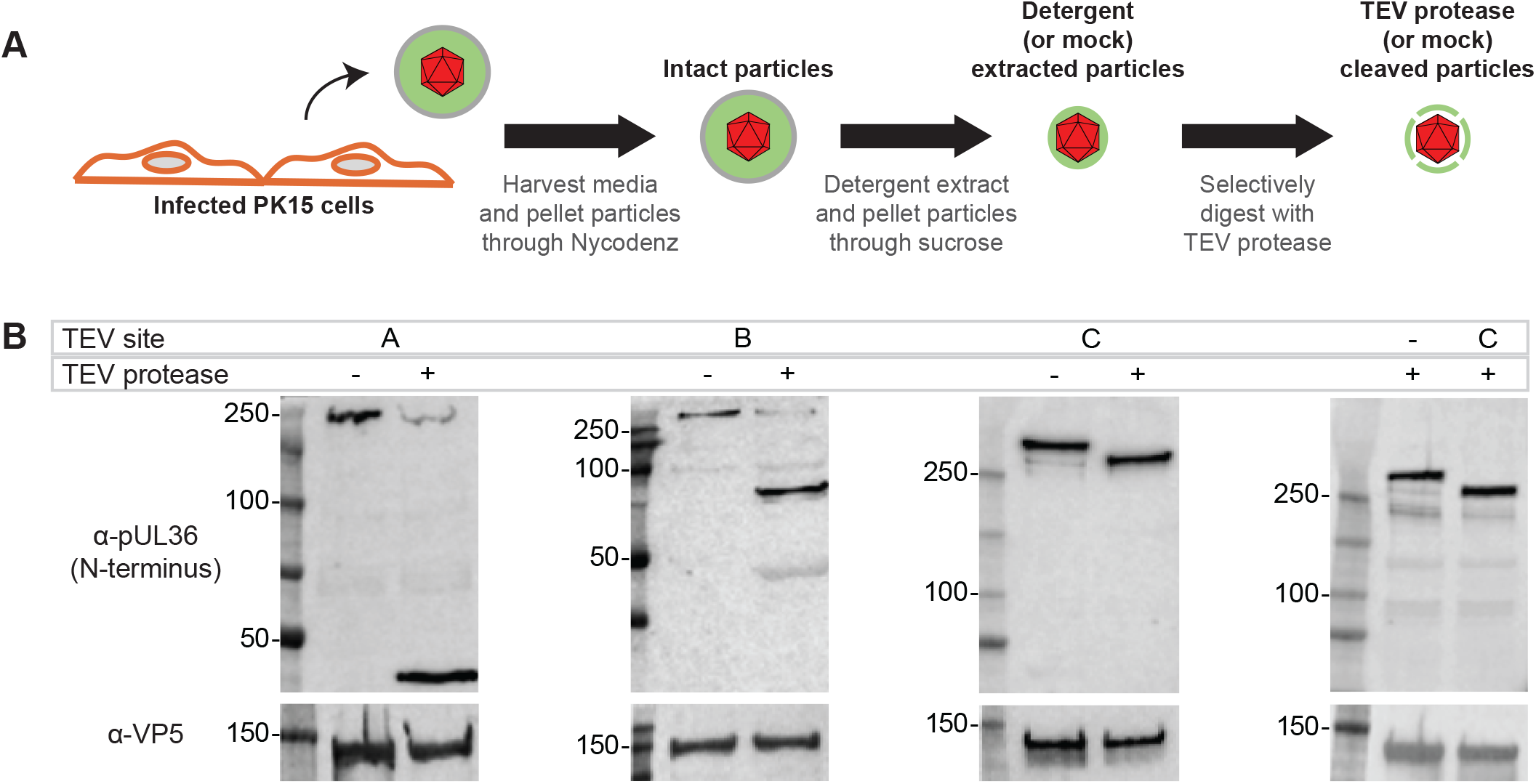
Recombinant TEV sites provide selective cleavage of pUL36. (A) Experimental workflow of extracellular virus particle harvesting and processing. (B) On-particle cleavage of pUL36 using viruses encoding a TEV site at position A, B, or C (see Figure 1A) was monitored by western blot analysis using an anti-pUL36 antibody that recognizes an N-terminal epitope and an anti-VP5 antibody as a loading control.

### Dissecting the capsid-pUL36 interface

To better understand how pUL36 contacts the capsid, a quantitative approach was developed that paired single-particle fluorescence imaging with TEV processing of dual-color-tagged viral particles. These experiments were designed to determine if pUL36 makes multiple contacts with capsids in extracellular viral particles (Fig. 1A) (18, 21, 24–26). Using a virus that contains a TEV site and two fluorophores (a mCherry-tagged capsid & GFP fused to the N-terminus of pUL36), the interface between pUL36 and capsids was probed. Specifically, if pUL36 contacts the capsid only through its C-terminus, then TEV cleavage should result in loss of GFP from particles; if instead pUL36 forms multiple contacts GFP should be retained. As a control, a virus encoding GFP fused to the C-terminus of pUL36 was included, which would not be expected to release GFP from the capsid following TEV protease treatment (18). A virus that lacked a TEV site but contained a tandem RFP-GFP fusion to the pUL35 hexon-tip capsid protein (RG-UL35) was included as an additional control (14). Capsids recovered from extracted extracellular viral particles were purified and reacted with TEV protease (Fig. 2A), which were then diluted and imaged to assay for retention of GFP fluorescence. In the absence of TEV, particles from all recombinant viruses displayed bright red- and green-fluorescence (14), and TEV treatment of the two control viruses did not reduce GFP emissions from the viral particles (Fig. 3). In contrast, particles containing a TEV site at position A or C in the GFP-pUL36 fusion emitted reduced green fluorescence following protease treatment. The loss of GFP emissions following cleavage at position C is noteworthy, as the result indicates that the three putative N-terminal capsid-binding sites are together insufficient for capsid binding under the conditions of the experiment. Given these results, the inability to release the GFP-UL36 N-terminus following cleavage at position B was unexpected. An *in silico* analysis of pUL36 with the ExPASy COILS software predicts that the region of pUL36 around site B is a hydrophobic coiled-coil. We suspect that cleavage at this site may expose hydrophobic residues because viral particles cleaved at position B had a tendency to form heterogeneous sized clumps consistent with aggregation. Regardless of the effect position-B cleavage incurred, cleavage at positions A and C demonstrated the efficacy of the TEV selective disassembly approach for studying viral particle architecture and revealed that pUL36 is stably anchored to capsids solely at its C-terminus.

**Figure 3:**
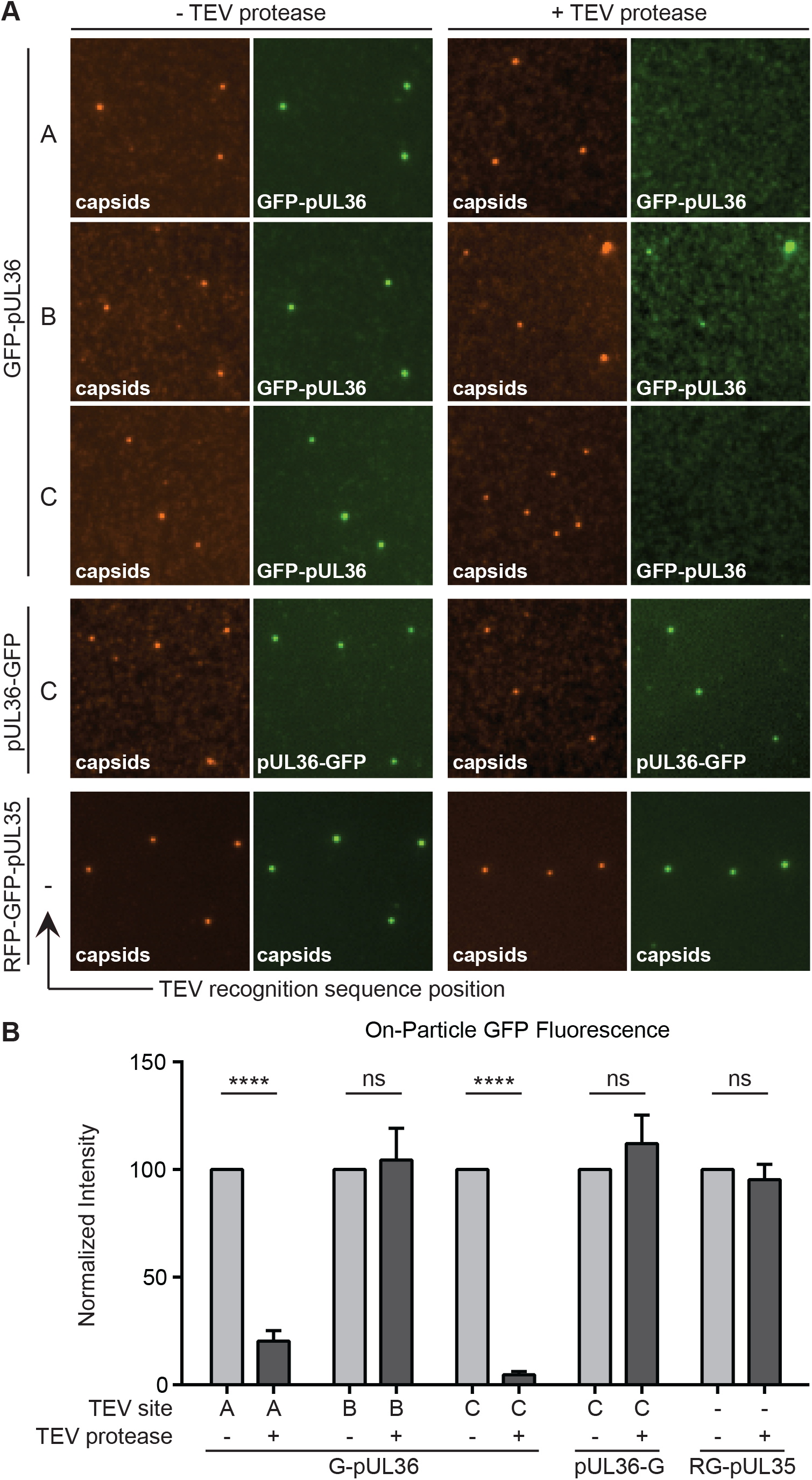
Mapping regions of pUL36 that maintain stable attachment to capsids. (A) Representative images of dual-fluorescent extracellular virus particles that were detergent extracted and incubated in the absence (left two columns) or presence (right two columns) of TEV protease. The position of the TEV site within pUL36 (position A, B, or C) is denoted, with a dash indicating no TEV site. All viruses encode the pUL25/mCherry capsid tag with the exception of the RFP-GFP-pUL35 dual-capsid tagged virus. (B) Fluorescence intensities from single particles represented in panel A. Values are normalized to the cognate ‘-TEV protease’ control and represent data from n ≥ 3 experiments. Error bars indicate standard error of the mean (**** = P < 0.001).

### Recombinant PRV strains for the analysis of additional tegument-capsid interactions

In addition to pUL36, other tegument proteins are proposed to interact with the capsid (16, 17, 51). Therefore, we extended the on-particle disassembly assay to determine whether six additional tegument proteins (pUS3, pUL16, pUL37, pUL47, pUL48, and pUL49) require pUL36 for stable capsid association in extracellular particles. A panel of dual-fluorescent viruses was constructed, each containing a TEV site within pUL36 at position C, an mCherry tag on the pUL25 capsid protein, and a GFP fusion to a tegument protein of interest. The position of the GFP fusion to each tegument protein was based on previous work, and analysis of released extracellular viral particles indicated that GFP-fused tegument proteins were incorporated at detectable levels in over 85% of particles (14). Recombinant viruses encoding GFP-tegument fusions grew to titers approximating wild-type PRV and had only moderately reduced propagation in PK15 cells relative to the parent non-GFP-tagged virus (Fig. 1C and Table 1).

### Differential susceptibility of tegument proteins to detergent

Many tegument proteins are involved in protein networks that connect the capsid to the membrane envelope (5). Because these interactions may be labile to detergent extraction, extracellular particles were exposed to 2% TX-100 to determine the degree to which each GFP-tegument fusion was solubilized from capsids on an individual particle level. Following purification from infected-cell supernatants viral particles were extracted and isolated by ultracentrifugation through a sucrose cushion. The particles were resuspended, spotted onto coverslips, and imaged by fluorescence microscopy as above (Fig. 2A). Representative images of mock- or detergent-extracted particles are presented in Figure 4A. A previously described image analysis algorithm was used to quantify red and green emissions from individual viral particles, which were first identified based on red-fluorescence intensities consistent with individual capsids (14). Representative GFP emission profiles in the presence and absence of detergent are shown for each virus in Figure 4B. In the absence of detergent, GFP emissions from each tegument fusion were variable from particle to particle such that the distribution of tegument incorporation was Gaussian, which was consistent with previous work examining tegument incorporation into individual PRV virions (14). Following detergent extraction, capsid-associated GFP emissions varied greatly depending on the tegument protein to which GFP was fused. The preservation of capsid-associated emissions from the GFP-pUL36, pUL37-GFP, and pUS3-GFP fusions indicated that the association of these proteins with capsids was resistant to the extraction conditions and furthermore that GFP fluorescence was not inactivated by the procedure. Affirming the latter point, red- and green-fluorescence signals from RFP-GFP-pUL35 particles were unchanged by detergent extraction (Fig. 4C), indicating that reductions of GFP signal observed with pUL16-GFP, pUL47-GFP, pUL48-GFP, and pUL49-GFP could be ascribed to protein loss from the particles (52). Although not critical for the subsequent analysis we note that following extraction, a fraction of either GFP-pUL36 or pUL37-GFP particles lost all detectable GFP emissions (25.5% and 21.7%, respectively; Figure 4B inset percentages). Whether this observation hints at a population of particles that lack pUL36-capsid interactions or is due to the labile nature of these proteins under some conditions will require further investigation (20, 21).

**Figure 4:**
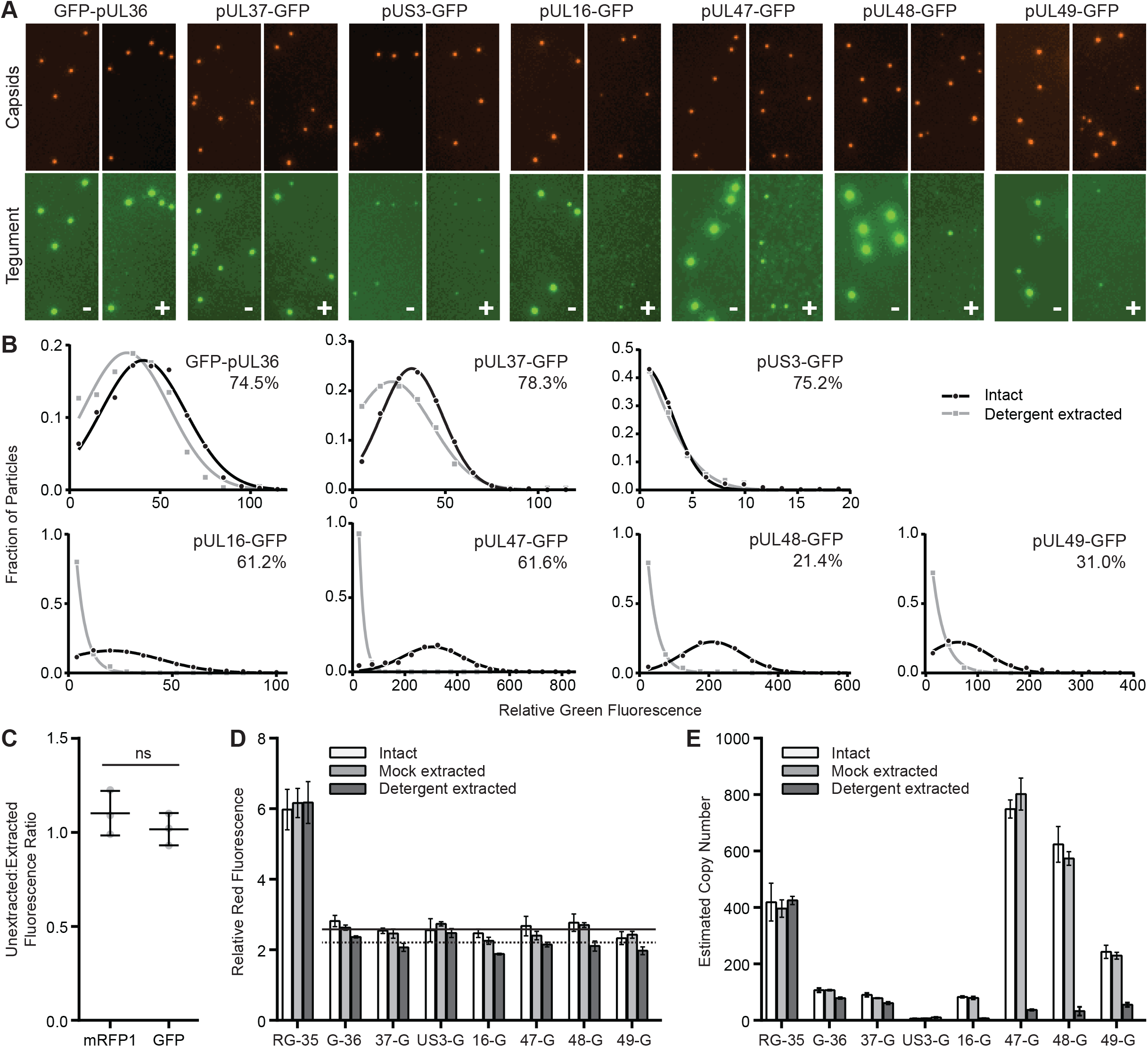
Tegument-capsid susceptibility to detergent extraction. (A) Dual-fluorescent extracellular viral particles containing the pUL25/mCherry capsid tag and GFP fused to a designated tegument protein were purified from the supernatant of infected cells and treated with buffer lacking (-) or containing (+) TX-100. Representative images of capsid and tegument emissions are shown, with each channel scaled equally across all samples. (B) GFP emission intensities from individual particles are show as histograms. The percentages in the upper right corner of each histogram indicate the proportion of particles with detectable GFP signal above threshold following detergent extraction. The data were fit to either Gaussian or decaying exponential distributions. All distributions had goodness of fit (R^2^) values > 0.95. (C) The red- and green-fluorescent signals of tandem-tagged mRFP-GFP-pUL35 particles were measured both before and after TX-100 extraction, and a -/+ TX-100 detergent ratio was calculated for each fluorescence channel (n=3). (D) Average red-capsid fluorescence intensities from purified particles following mock or TX-100 extraction (n=3 per condition). Error bars indicate standard error of the mean. The solid horizontal line is the average intensity (“Intact” + “Mock extracted”) across the seven capsid-tegument dual-fluorescent viruses. The dotted horizontal line is the average intensity following detergent extraction. (E) Estimated protein copy numbers based on average green-fluorescence intensities (see Methods). Error bars indicate standard error of the mean.

Fluorescence intensity measurements made from three independent experiments per virus were compiled to extrapolate estimated protein copy numbers before and after detergent extraction (Fig. 4D and E). As a technical point, these results demonstrated that the secondary ultracentrifugation step through a sucrose cushion did not alter red- or green-fluorescent intensities as compared with virions isolated by a single purification step alone (compare “Intact” and “Mock-extracted” samples). However, detergent extraction was noted to consistently produce a slight reduction in red-fluorescence from the pUL25/mCherry fusion encoded by any of the seven capsid/tegument dual-tagged viruses. The red-fluorescence intensity measurements in the seven mock-extracted samples were averaged (Fig. 4D, solid horizontal line), and this intensity value was assigned a copy number of 70.5 based on the pUL25 copy number reported in a previously study (14). By comparing the mock extracted average red-fluorescent intensity to a similar average across the detergent extracted condition (Fig. 4D, dashed horizontal line), we extrapolated that the pUL25/mCherry copy number was reduced to 60.2 ± 5.4 following exposure to detergent. Interestingly, this number is consistent with five copies of pUL25/mCherry at each of the twelve capsid vertices (53, 54). While this value could be coincidental, it is possible that 60 copies of the pUL25/ mCherry fusion are stably bound to capsids around the vertices, while a smaller number of pUL25 copies incorporate into virions by another means. We also note that a recent cryoEM reconstruction of PRV reveals that there are 10 copies of pUL25 per vertex, further indicating that fusing mCherry to pUL25 may reduce its copy number on capsids (6). Green-fluorescence intensity measurements were converted to copy numbers by normalizing the average intensities to the reported copy number for GFP-pUL36, which is 107.1 copies/virion (Fig. 4E and Table 3) (14). For several of the tegument-tagged viruses, GFP emissions were significantly reduced in both overall intensity/particle and in terms of the number of particles with GFP signal above threshold following detergent extraction (Fig. 4B and E; Table 3). This precluded the inclusion of the dual-fluorescent TEV-site containing viruses encoding GFP fusions to pUL16, pUL47, pUL48, pUL49 in the subsequent TEV protease disassembly assay.

**Table 3:** Protein copies per extracellular viral particle

| | Copy # (± SD) | |
| --- | --- | --- |
|  | Unextracted | Extracted |
| GFP-pUL36 | 107.1<br>(1.4) | 79.1<br>(2.2) |
| pUL37-GFP | 78.8<br>(0.9) | 61.1<br>(3.5) |
| pUS3-GFP | 6.7<br>(0.1) | 10.3<br>(1.6) |
| pUL16-GFP | 79.6<br>(4.2) | 7.4<br>(0.5) |
| pUL47-GFP | 802.0<br>(46.6) | 37.0<br>(2.2) |
| pUL48-GFP | 574.0<br>(19.8) | 32.6<br>(12.5) |
| pUL49-GFP | 229.4<br>(9.8) | 55.7<br>(6.1) |

### pUS3 and pUL37 connectivity to capsids in extracellular particles occurs through pUL36

The nature of the attachment of pUS3 and pUL37 to capsids was further explored by testing if these proteins were tethered to capsids via pUL36. Viral particles were isolated from the supernatant of cells infected with dual-fluorescent viruses encoding a TEV site at position C within pUL36, extracted, and incubated in the presence or absence of TEV protease (Fig. 5). Fluorescent intensities from individual viral particles were measured as described above. TEV protease treatment resulted in significant loss of both proteins from capsids, indicating that the pUS3 and pUL37 tegument proteins are bound to capsids indirectly via pUL36 and make no additional stable contacts with capsids.

**Figure 5:**
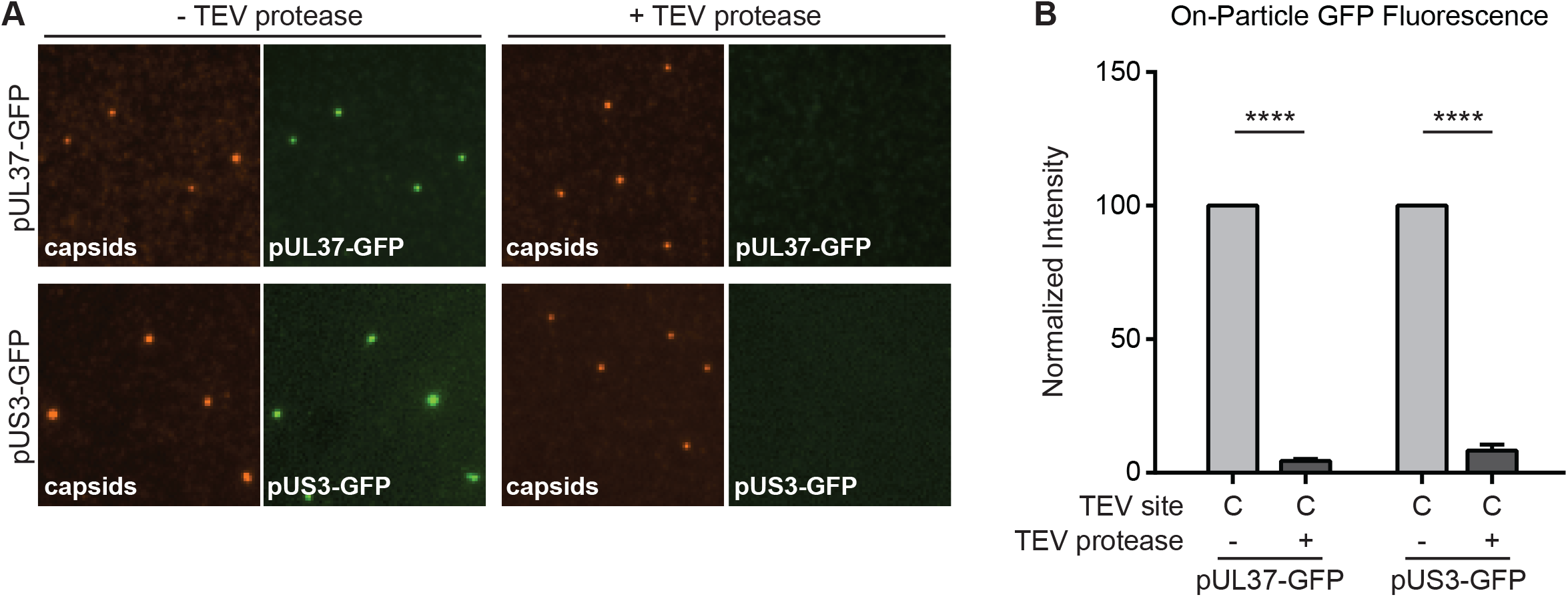
The pUS3 and pUL37 tegument proteins are tethered to capsids by pUL36. (A) Extracted dual-fluorescent extracellular virus particles encoding a TEV site at position C in pUL36 were incubated in the absence or presence of TEV protease, and representative images of fluorescence emissions are shown. (B) Fluorescent intensities from single particles are presented normalized to the cognate ‘-TEV protease’ control (n≥3). Error bars indicate standard error of the mean (**** = P < 0.001).

## DISCUSSION

The herpesvirus tegument is notable both for its contribution to particle architecture, as well as for effector functions that commence immediately upon entry into cells and subsequently act throughout the infectious cycle (5). Whereas tegument proteins are targeted for assembly into nascent virions late in the infectious cycle, upon initial infection they selectively disassemble such that only a subset remain capsid bound during cytoplasmic transit and nuclear delivery (38, 55–57). These properties attest to the dynamic nature of the tegument, which is further highlighted by the finding that at least one tegument protein reconfigures within the intact virion upon cell contact (51). Alphaherpesviruses incorporate > 20 types of tegument proteins (12, 13), some represented in hundreds of copies, with the total tegument mass constituting approximately half of the virion. While many pairwise interactions between viral tegument proteins have been described (5), it is not clear how these proteins coalesce in a mature particle. In this study, we developed a selective disassembly assay based on the TEV site-specific protease to investigate how several tegument proteins anchor onto the capsid in extracellular virus particles. Whereas traditional mutagenesis approaches have advanced our understanding of virion morphogenesis, this new approach allowed for the targeted dissection of the assembled viral particle. In this initial application of the method, we identified that PRV pUL36 is stably bound to capsids solely by its C-terminus, and that tegument proteins pUS3 and pUL37 make no additional stable contacts with the capsid but are instead dependent upon pUL36 for stable tethering to the capsid (Fig. 6).

**Figure 6:**
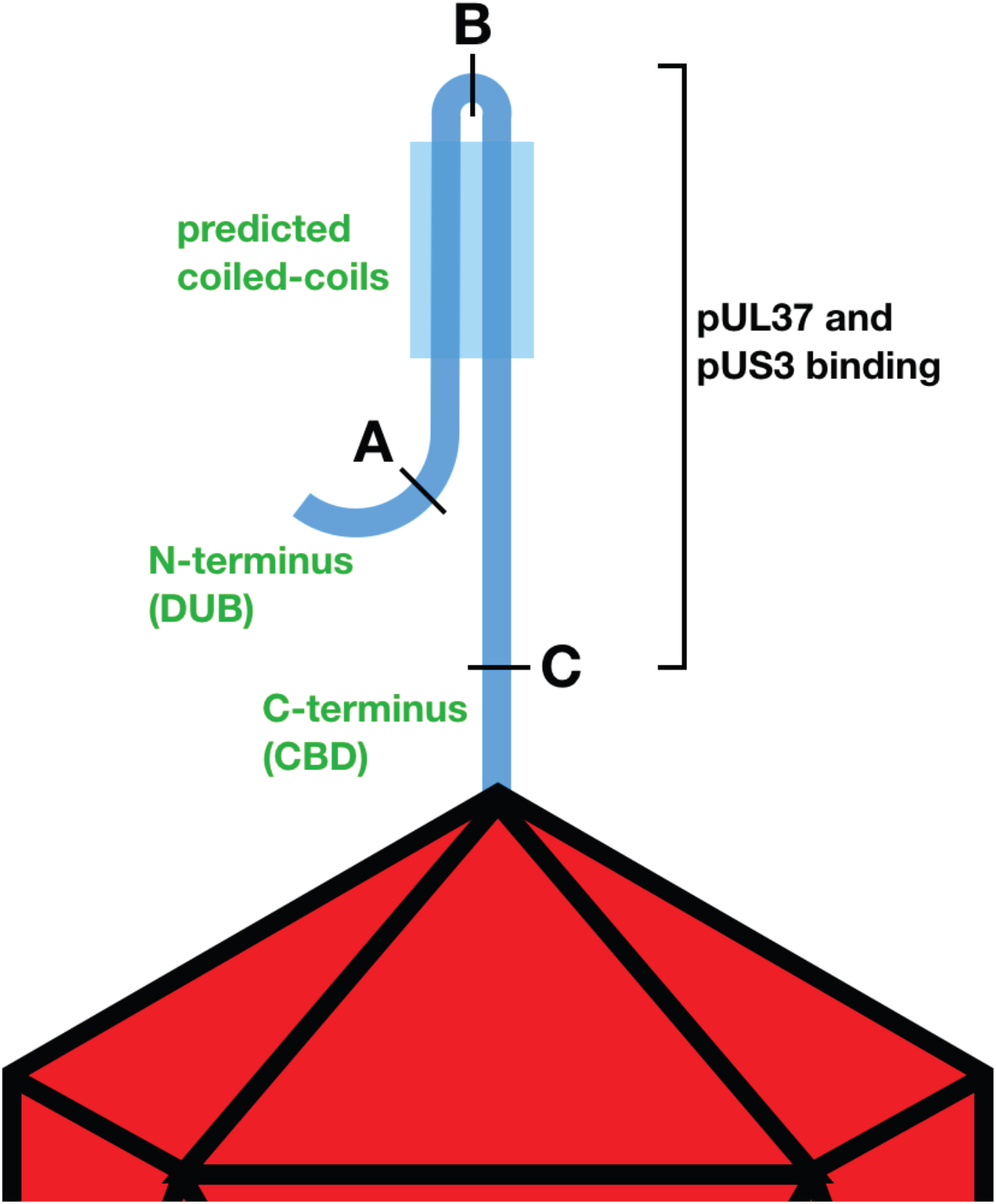
Model of pUL36 topology on capsids from extracellular particles. Simplified illustration of a single copy of pUL36 shown bound to a vertex of the herpesvirus capsid by its C-terminal capsid-binding domain (CBD). The N-terminus of pUL36 up to position A, housing the deubiquitinase catalytic domain (DUB), makes no stable contacts with the remainder of pUL36 or the capsid surface. Position B is located in a region of predicted coiled-coils and cleavage at this site does not release the N-terminus of pUL36 from the capsid. Cleavage at position C releases the amino terminal peptide (which constitutes the bulk of the pUL36 protein) from the capsid, indicating that only the sequence C-terminal of position C affords stable attachment to capsids under the conditions of these experiments. Two capsid-associated tegument proteins, pUL37 and pUS3, are also released upon cleavage at position C, indicating that these proteins are dependent upon pUL36 for stable capsid association.

We previously used quantitative fluorescence imaging to localize tegument protein populations within extracellular PRV particles, resulting in an intravirion map of tegument distributions (14). That prior study also provided copy numbers of GFP-tagged tegument proteins and their variance from particle to particle. Building on that quantitative imaging approach, we investigated how tegument proteins are affected by detergent extraction. We note that this study differed from our prior analysis in that capsids were tagged with a pUL25/mCherry fusion, which better preserves the properties of wild-type virus than the earlier mRFP1-VP26 capsid reporter (36). Nevertheless, the estimated copy numbers of GFP-tegument fusion proteins in intact viral particles was on par between the studies, with the exception of pUL47 and pUL48, which were present in higher copies in the current report. Four tegument proteins (pUL16, pUL47, pUL48, and pUL49) were susceptible to extraction in 2% TX-100 and 0.5 M NaCl and were not retained uniformly on the capsid surface following solubilization. The histogram profiles of these proteins following extraction (i.e. decaying exponentials) suggest that the remaining residual proteins was retained in a non-specific or stochastic fashion. The loss of pUL48 is notable as this tegument protein interacts with pUL36 in HSV-1 to promote particle assembly (58–60). This finding highlights that interactions important for virion assembly are not necessarily preserved in mature particles, which is the case for the pUL16 tegument protein (16, 51). Three tegument proteins (pUL36, pUL37, and pUS3) were significantly retained on detergent-extracted particles, and the emission profiles from these GFP-fusions remained Gaussian, indicating they remained capsid bound (14). The results from the present study are largely consistent with prior studies that have examined differential extraction of bulk populations of extracellular herpesvirus particles using salt, nonionic detergent, reducing agents, or phosphorylation (20, 61–65). The differences observed in tegument extractability between studies indicate that the isolation, handling, and extraction conditions effect experimental outcomes. Nevertheless, these results are consistent with pUL36, pUL37, and pUS3 remaining capsid bound upon entering cells (38, 55, 56, 66, 67). We infer that the retention of these proteins on capsids upon cell entry is due to their collective attachment to the capsid surface by the C-terminus of pUL36.

Despite these findings, several prior studies have implicated the presence of capsid binding sites in pUL36 in addition to the C-terminal capsid-binding domain (21, 24–26). Furthermore, pUL36 can assemble into extracellular particles when the C-terminal capsid-binding site is eliminated, although it is subsequently lost from capsids upon entry into cells (68). This indicates that pUL36 assembly into virions can occur by a mechanism that is distinct from its ability to form a stable attachment to the capsid surface, likely by the pUL48 tegument protein as evidenced by its role in recruiting pUL36 into extracellular light particles that lack capsids (69). However, recruitment by pUL48 would not explain how the capsid is brought into the nascent forming virion in the absence of the UL36 C-terminus. While more research will be needed to parse out how the tegument layer attaches to capsids, it seems that pUL36 either makes additional contacts that are not stable or there is another tegument protein apart from those included in this study that serves as a link between the capsid and tegument. The bulk of the tegument is surprisingly plastic in its composition and many tegument proteins are individually dispensable for virion assembly. While we cannot rule out that there may be redundant tegument-capsid interactions that promote particle assembly, cryoEM reconstruction of PRV and HSV-1 virions do no attest to capsid interactions apart from pUL36 (6, 7, 70).

This study provides new details about the alphaherpesvirus tegument network. The selective disassembly assay developed for these studies should prove useful for future investigations of the viral particle architecture and its dynamics, including deciphering changes in the tegument that can occur temporally or following heparin cross-linking of glycoprotein C (16, 63). Because diverse herpesviruses share many architectural features and a common particle morphology, insights from this study may be informative to viruses across the *herpesviridae* family.

## ACKNOWLEDGEMENTS

We thank Sarah Antinone for helping edit the manuscript. The monoclonal antibody directed against the PRV VP5 protein was generously provided by Lynn Enquist. Sequencing services were performed at the Northwestern University Genomics Core Facility.

